# Neural mechanisms of impaired self-compassion in depressed adolescents during social exclusion

**DOI:** 10.64898/2026.08.04.741714

**Authors:** Huijing Sun, Wenjing Zou, Duanwei Wang, Yihao Zhang, Changlin Bai, Wenfu Li, Yanan Xiao, Xiuqing Niu, Xuexiao Shao, Xiaoyue Wang, Bernhard Hommel, Ying Yang, Kangcheng Wang

**Affiliations:** School of Psychology, Shandong Provincial Key Laboratory of Brain Science and Mental Health, Shandong Normal University, Jinan, 250358, China; Shandong Mental Health Center, Shandong University, Jinan, 250014, China; School of Rehabilitation Medicine, Jining Medical University, Jining, 272067, China; Institute of Brain Science and Brain-inspired Research, Shandong First Medical University & Shandong Academy of Medical Sciences, Jinan, 250117, China; Cheeloo College of Medicine, School of Basic Medical Sciences, Shandong University, Jinan, 250100, China

**Keywords:** Adolescents with depression, social exclusion, self-compassion, EEG

## Abstract

**Background:** Self-compassion has shown to be an effective emotion regulation strategy, but its neural mechanisms remain understudied in adolescents with depression who suffer from social exclusion. This study employed event-related potentials to investigate the psychophysiological mechanisms underlying impaired self-compassion in adolescents with depression during social exclusion.

**Methods:** Thirty-seven adolescents with major depressive disorder (age = 16.46 ± 2.02) and thirty-two demographically matched healthy controls (age = 16.54 ± 2.29) completed a social exclusion scenario imagination task. This task presented participants with social exclusion scenarios, asking them to imagine themselves as the excluded individual and subsequently rate their negative emotions. The regulation session required participants to use self-compassionate statements to regulate their emotional responses while imagining and rate how much self-compassion they have engaged in; the non-regulation required them not to use those statements and rate their negative emotions immediately following imagination. EEG signals were recorded during the task, and the late positive potential (LPP) was analyzed to examine the neural responses associated with self-compassion regulation following social exclusion.

**Results:** Compared with the non-regulation condition, both groups showed less negative emotion during the self-compassion regulation condition. Adolescents with depression exhibited significantly lower self-compassion ratings than healthy controls, and showed a significantly smaller decrease in negative emotions. Higher self-compassion ratings were correlated with greater emotion regulation effect, particularly in adolescents with depression. Furthermore, LPP amplitudes were significantly higher in adolescents with depression than in healthy controls during both regulation and non-regulation conditions.

**Conclusions:** Adolescents with depression were characterized by impairment in using self-compassion to downregulate negative emotions when confronted with social exclusion. LPP amplitudes were consistently elevated in adolescents with depression, underscoring their potential as a critical focus for psychophysiologically-informed therapies.

## 1. Introduction

Adolescence is a turbulent transitional period characterized by profound physiological and psychological changes [1, 2]. These rapid changes make adolescents especially vulnerable to mood disorders [3–5]. Depression, a common and increasingly prevalent mood disorder, is associated with significant cognitive and affective impairments [6–8]. Importantly, adolescents with depression often exhibit negative attribution biases, along with atypical emotional responsivity and regulation [9–11]. These emotional difficulties are particularly pronounced in socially adverse contexts, such as social exclusion, where adolescents with depression show prominent deficits in emotional processing and regulation [12–14]. Additionally, these individuals show dysfunctions in the executive control network, which reflects impaired control ability of emotional salience processing and regulation during negative social interactions [12, 15]. Consequently, adolescents with depression face greater difficulty in coping with the arising negative emotions caused by adverse social experiences relative to healthy adolescents [16–18]. Consistent with these findings, studies of adolescents in interpersonal contexts have consistently documented potentiated negative reactivity, with depressive symptoms amplifying emotional responses to daily peer rejection and to experimentally induced interpersonal rejection [16, 17]. Because emotional reactivity increases during adolescence while prefrontal regulatory capacities are still maturing, and because peer evaluation is especially salient at this stage, adolescents with depression may be particularly prone to emotion regulation difficulties when facing interpersonal stress. Identifying effective strategies to alleviate such emotional disturbances is therefore crucial, as it may help mitigate the adverse impact of depression on adolescents.

Social exclusion is a broad concept encompassing various adverse interpersonal experiences, including emotional, physical or other forms of isolation from others, as well as experience of social devaluation [19–21]. Unlike ostracism (being ignored without explanation) or rejection (explicit communication that one is unwanted), social exclusion encompasses both of these specific forms and a wider range of exclusionary experiences, such as being left out of peer activities, being ignored during social interactions, or perceiving a lack of acceptance from peers [20, 22]. Experiencing social exclusion threatens psychological needs and often results in decreased self-esteem, increased psychological distress, and impaired well-being [20–22]. These negative consequences are particularly pronounced in adolescents with depression, who already face heightened vulnerability to social stressors [13, 16]. This is consistent with the negative potentiation account [23], which posits that social exclusion may elicit amplified negative emotional responses in adolescents with depression. Such amplified reactivity underscores the need for targeted emotion regulation strategies to mitigate the harmful emotional impact of social exclusion. By fostering self-acceptance rather than self-criticism in response to social adversity, self-compassion may help alleviate the negative impact of social exclusion among adolescents with depression [24].

Self-compassion is a positive self-attitude characterized by kindness toward oneself, particularly in the face of failures, mistakes or difficulties [25, 26]. It includes three core components: self-kindness (versus self-judgment), common humanity (versus isolation), and mindfulness (versus over-identification) [26, 27]. For instance, when adolescents experience emotional distress following being excluded during a peer interaction, they may acknowledge that rejection is a common aspect of adolescent social development and that these emotions are temporary. Such self-compassion strategies would reduce their self-criticism and rumination, thereby contributing to improved mental health outcomes [25, 26]. Similarly, studies in adolescent populations indicate that higher levels of self-compassion are strongly associated with lower levels of psychological distress [28]. This evidence has suggested self-compassion functions as an adaptive emotion regulation strategy to relieve negative emotions and enhance mental well-being [29–32]. However, compared with never-depressed individuals, depressed adults reported less self-compassion, which was negatively associated with symptom-focused rumination [33]. By now, the neural mechanisms underlying impaired self-compassion in adolescents with depression particularly under social exclusion remain poorly understood.

Neuroimaging studies provide valuable insights into the neural correlates of self-compassion, while event-related potentials (ERPs) offer complementary advantages by capturing the temporal dynamics of emotional processing and regulation with millisecond precision. The late positive potential (LPP), a sustained occipital-parietal ERP component, has been established as a reliable neural marker of emotional processing and regulation [34–37]. LPP amplitude reflects sustained attention and elaborated processing of emotionally salient stimuli, with greater amplitude indicating deeper emotional engagement [37, 38]. Critically, LPP demonstrates sensitivity to emotion regulation strategies: successful regulation typically reduces LPP amplitude, reflecting diminished emotional reactivity to negative stimuli [38]. Individuals with depression exhibit enhanced processing of self-relevant stimuli, as well as amplified LPP amplitudes in response to negative affective stimuli [39]. Therefore, the LPP may provide a promising neural indicator for investigating self-compassion regulation mechanisms in adolescents with depression, particularly in contexts of social distress such as exclusion. However, few studies have directly examined the electrophysiological correlates of self-compassion regulation.

The current study aimed to examine the effects of self-compassion regulation in depressed versus healthy adolescents at both behavioral and electrophysiological levels. We recruited adolescent patients with depression and healthy controls to perform a scenario imagination task presenting social exclusion scenarios. Participants were instructed to imagine themselves as the excluded individual and, depending on the experimental condition, either apply or not apply self-compassionate statements to comfort themselves while imagining exclusion scenarios. ERPs were recorded while participants were completing the task, with the LPP as the focus of our analysis. We hypothesized that (1) self-compassion would reduce negative emotions in both adolescents with depression and healthy controls; (2) adolescents with depression would engage less self-compassion and exhibit elevated LPP amplitudes relative to healthy adolescents during social exclusion.

## 2. Methods

### 2.1. Participants

A total of 37 adolescent patients with depression were recruited from Shandong Mental Health Center (age = 16.46 ± 2.02, 12–20 years, male/female = 8/29; Table 1). Inclusion criteria for the depressed group were: (a) a diagnosis of major depressive disorder according to DSM-5 criteria, confirmed by two certified psychiatrists; (b) a score > 15 on the Children’s Depression Inventory (CDI) [40]; (c) being able to comprehend the questionnaires and complete the experimental procedures. Exclusion criteria included: (a) comorbid psychiatric disorders such as schizophrenia, obsessive-compulsive disorder, bipolar disorder, or substance use disorder; (b) neurological disorders or serious physical illnesses; (c) secondary depression caused by organic diseases or medication use.

**Table 1.** Demographic and clinical characteristics of adolescents with depression and healthy controls.

| Variables | All participants<br>(n=69) | Healthy Controls<br>(n=32) | Depressed adolescents<br>(n=37) | $t/\chi^2$ | $p$ |
| --- | --- | --- | --- | --- | --- |
| Age, years | 16.50(2.13) | 16.54(2.29) | 16.46(2.02) | -0.169 | 0.867 |
| Sex, n (%) |  |  |  | 0.119 | 0.730 |
| Male | 17(24.60) | 9(28.10) | 8(21.62) |  |  |
| Female | 52(75.40) | 23(71.90) | 29(78.38) |  |  |
| Education, n (%) |  |  |  | 2.309 | 0.315 |
| ≤6 years | 4(5.80) | 3(9.37) | 1(2.70) |  |  |
| 7-9 years | 22(31.88) | 8(25.00) | 14(37.84) |  |  |
| ≥10 years | 43(62.32) | 21(65.63) | 22(59.46) |  |  |
| Parental marital status, n (%) |  |  |  |  |  |
| Married | 66(95.65) | 32(100.00) | 34(91.89) | 1.113 | 0.291 |
| Divorced/other | 3(4.35) | 0(0.00) | 3(8.11) |  |  |
| Single child families, n (%) |  |  |  |  |  |
| Yes | 19(27.54) | 9(28.13) | 10(27.03) | 0.000 | 1.000 |
| No | 50(72.46) | 23(71.88) | 27(72.97) |  |  |
| Family psychiatric history, n (%) |  |  |  |  |  |
| Yes | 2(2.90) | 0(0.00) | 2(5.41) | 0.378 | 0.538 |
| No | 67(97.10) | 32(100.00) | 35(94.60) |  |  |
| CDI total score | 18.48(8.90) | 12.31(5.61) | 23.81(10.97) | -5.588 | <0.001 |
| Negative self-esteem | 3.38(1.93) | 2.25(1.22) | 4.35(2.38) | -4.701 | <0.001 |
| Negative mood | 3.78(2.52) | 1.97(1.56) | 5.35(3.12) | -5.813 | <0.001 |
| Anhedonia | 5.41(3.20) | 3.53(2.69) | 7.03(3.58) | -4.529 | <0.001 |
| Interpersonal problem | 1.75(1.40) | 1.28(0.99) | 2.16(1.68) | -2.699 | 0.009 |
| Ineffectiveness | 4.16(1.89) | 3.28(1.55) | 4.92(2.14) | -3.590 | <0.001 |
| MASC total score | 49.39(21.16) | 38.06(14.40) | 59.19(21.32) | -4.877 | <0.001 |
| Physical symptoms | 12.78(9.51) | 6.31(4.17) | 18.38(9.30) | -7.108 | <0.001 |
| Harm avoidance | 14.72(4.87) | 13.53(4.64) | 15.76(4.89) | -1.930 | 0.058 |
| Social anxiety | 14.91(6.61) | 12.91(6.22) | 16.65(6.53) | -2.426 | 0.018 |
| Separation anxiety | 6.43(4.96) | 4.88(3.81) | 7.78(5.47) | -2.523 | 0.014 |

Another 32 age- and sex-matched healthy adolescents were recruited through public advertisements (age = 16.54 ± 2.29, 11–20 years, male/female = 8/24; Table 1). There were no significant group differences in terms of sex (*χ*^2^ = 0.110, *p* = 0.740) or age (*t* = 0.169, *p* = 0.867). Inclusion criteria for healthy controls were: (a) no current or historical psychiatric disorder based on DSM-5 criteria; (b) no history of psychiatric illnesses in first-degree relatives. Exclusion criteria included: (a) neurological conditions or serious physical illness; (b) a history of substance abuse or dependence.

Demographic characteristics, including age, sex, education level, parental marital status, single-child family status, and family psychiatric history, were collected and compared between groups. No significant group differences were observed in any of these sociodemographic variables (all *ps* > 0.05; Table 1). Depressive symptoms were assessed using the CDI and anxiety symptoms were assessed using the Multidimensional Anxiety Scale for Children (MASC) [41]. As expected, adolescents with depression scored significantly higher than healthy controls on both the CDI (*t* = -5.59, *p* < 0.001) and the MASC (*t* = -4.88, *p* < 0.001; Table 1).

This study was approved by the Ethics Committee of Shandong Normal University and Shandong Mental Health Center and conducted in accordance with the Declaration of Helsinki. Written informed consent was obtained from all participants and their legal guardians. Participants received monetary compensation upon completing the study.

### 2.2. The scenario imagination task

We utilized a “self-compassion regulation under social exclusion” task to assess self-compassion in adolescents [42]. The task consisted of two experimental conditions: one involving self-compassion regulation and the other without it. Prior to the experiment, all participants were instructed to write down three compassionate sentences. For example, a self-compassionate sentence generated by a participant was: “Everyone will experience the moment of rejection, which does not mean that I am not worthy of love.” Previous studies have demonstrated that such self-compassionate statements can efficiently activate a compassionate mindset [43, 44].

This scenario imagination task consisted of two blocks, with 30 trials per block. Each trial began with a fixation cross (random time, 1–3 seconds) presented at the center of the screen, followed by an image depicting a social exclusion scenario for 8 seconds. All exclusion-related images were selected from the Image Database of Social Inclusion and Exclusion in Young Asian Adults [45], which have been validated in prior work [42]. The same 30 images were used in both blocks, with trials pseudo-randomized in each block. In Block A, participants were instructed to imagine themselves as the excluded individual in each image without thinking of the self-compassion statements they had generated prior to the experiment. After viewing each image, participants rated their negative emotion experience on a 9-point scale (1 = *not at all*, 9 = *extremely sad*). In Block B, participants were instructed to imagine themselves as the excluded individual and actively apply self-compassion statements to comfort themselves. After viewing each image, participants were firstly asked to rate the extent to which they engaged in self-compassion on a 9-point scale (1 = *none*, 9 = *extremely engaged*), followed by a rating of their negative emotional experience. The order of the two blocks was counterbalanced across participants to control for potential order effects (Figure 1).

**Figure 1.** Experimental paradigm for imagining social exclusion with and without self-compassion regulation. The task consisted of 60 trials, with each participant completing Block A and Block B (30 trials each). (A) Participants were instructed to imagine themselves as the adolescent circled in red and to imagine and feel according to the task requirements for 8 seconds, after which they rated their negative emotion with 9-point scale (1 = not at all, 9 = extremely sad). (B) In the regulation condition, participants additionally engaged in self-compassion after viewing the image and then rated both their self-compassion (1 = none, 9 = extremely engage) and negative emotion. Block order was counterbalanced across participants.

### 2.3. EEG recording and preprocessing

Electroencephalography (EEG) data were recorded throughout the task to examine the neural activity in the context of social exclusion and self-compassion regulation. EEG data were recorded using a 64-channel Quik-Cap (Compumedics Neuroscan, Charlotte, NC, USA) in accordance with the extended 10-20 system. Two additional electrodes were placed above and below the left eye to record vertical electrooculogram activity associated with blinks and vertical eye movements. To ensure high-quality signal acquisition, electrode impedances were maintained below 10 kΩ throughout recording. EEG signals were amplified using a SynAmps 2 amplifier (Compumedics Neuroscan, Charlotte, NC, USA) and the sampling rate was 1000 Hz. Before EEG recording, participants were instructed to minimize head and body movements, minimize unnecessary eye movements, and maintain a relaxed posture throughout the experiment. Participants were not asked to repeat the task if there was excessive movement because repeating experimental blocks could introduce additional practice and habituation effects. During recording, EEG signal quality and electrode impedances were continuously monitored to ensure stable data acquisition.

Offline EEG preprocessing was performed following standard ERP analysis procedures. EEG signals were first filtered using a band-pass filter of 0.1–30 Hz. Bad channels were identified based on abnormal signal characteristics and interpolated using spherical spline interpolation based on the original channel locations. The continuous EEG data were then segmented into epochs ranging from -200 to 1100 ms time-locked to the onset of social exclusion-related images. Independent component analysis (ICA) was performed to identify and remove ocular and other non-neural artifacts based on component topographies and time courses. Epochs containing residual artifacts, including excessive voltage fluctuations exceeding ±80 μV or severe baseline drift, were rejected before ERP averaging. Baseline correction was applied using the -200 to 0 ms pre-stimulus interval.

To ensure reliable ERP estimation, participants with fewer than 20 valid trials in either condition or with excessive EEG artifacts were excluded from further analyses [46]. A total of 18 participants were excluded due to EEG quality issues, including 3 participants with missing EEG data or recording errors, 7 participants with excessive artifacts (e.g., head movements), and 8 participants with more than 10 bad channels. After EEG preprocessing and quality control, a total of 51 participants were included in the final ERP analysis, including 25 in the depressed group (age = 16.22 ± 2.10, male/female: 5/20) and 26 in the healthy control group (age = 16.66 ± 2.36, male/female: 6/20). Analyses were conducted to examine whether EEG-related exclusion introduced potential selection bias. Demographic and clinical characteristics were compared between participants being retained in the final ERP analysis (*n* = 51) and those being excluded due to EEG quality issues (*n* = 18). There were no significant group differences in sex distribution (*χ*^2^ = 0.74, *p* = 0.389), age (*t* = 0.34, *p* = 0.735), CDI total scores (*t* = -0.38, *p* = 0.710), or MASC total scores (*t* = -0.18, *p* = 0.858). These findings indicated that EEG artifact-related exclusion did not introduce systematic bias into the final ERP sample. Following artifact correction, clean epochs were averaged separately for each participant and experimental condition to generate ERP waveforms. The late LPP was quantified as the mean amplitude within the 600–1100 ms time window following stimulus onset, consistent with previous ERP studies examining sustained emotional processing [47]. Based on previous studies [48], PO3, PO4 and Poz electrodes were selected for LPP analysis, and the LPP amplitude was calculated by averaging signals across these three electrode sites.

### 2.4. Statistical analysis

All analyses were conducted using R software (4.4.2). Independent samples t-test was used to examine group differences in self-compassion ratings, and negative emotion ratings under the two experimental conditions. Paired t-tests were conducted to assess the effects of self-compassion regulation on negative emotion ratings within each group. The emotion regulation effect was quantified as the magnitude of reduction in negative emotion ratings between the non-regulation and self-compassion regulation conditions and was compared between groups. Partial correlation analyses examined relationships between self-compassion ratings and the emotion regulation effect, with age and sex included as covariates. A stratified linear regression further probed the predictive roles of self-compassion ratings, group assignment, and their interaction on the emotion regulation effect.

EEG data were analyzed by comparing LPP amplitudes. Independent samples t-tests compared groups under each experimental condition. A repeated measures analysis of variance (ANOVA) was performed to identify main and interaction effects of group (healthy controls vs. depressed patients) and condition (self-compassion regulation vs. non-regulation) on LPP amplitude.

## 3. Results

### 3.1. Behavioral effects of self-compassion regulation

Adolescents with depression showed significantly lower self-compassion ratings than healthy controls (*t* = 3.43, *p* < 0.001, Figure 2A). In the non-regulation condition, the two groups did not differ significantly in negative emotion scores (*t* = 0.57, *p* = 0.570); in the self-compassion regulation condition, the depressed group reported greater negative emotion than healthy controls (*t* = 2.54, *p* = 0.013, Figure 2B). Self-compassion regulation significantly reduced negative emotion ratings both in the depressed group and healthy controls (Figure 2C). Moreover, the reduction in negative emotion between regulation and non-regulation conditions was significantly smaller in the depressed group (*t* = 2.53, *p* = 0.014, Figure 2D), indicating a smaller emotion regulation effect in adolescents with depression. Higher self-compassion ratings were correlated with greater emotion regulation effects across both groups (*r* = 0.306, *p* = 0.011; Figure 3). This association was significant within the depressed group (*r* = 0.331, *p* = 0.045), but not in healthy controls (*r* = 0.165, *p* = 0.367).

**Figure 2.**
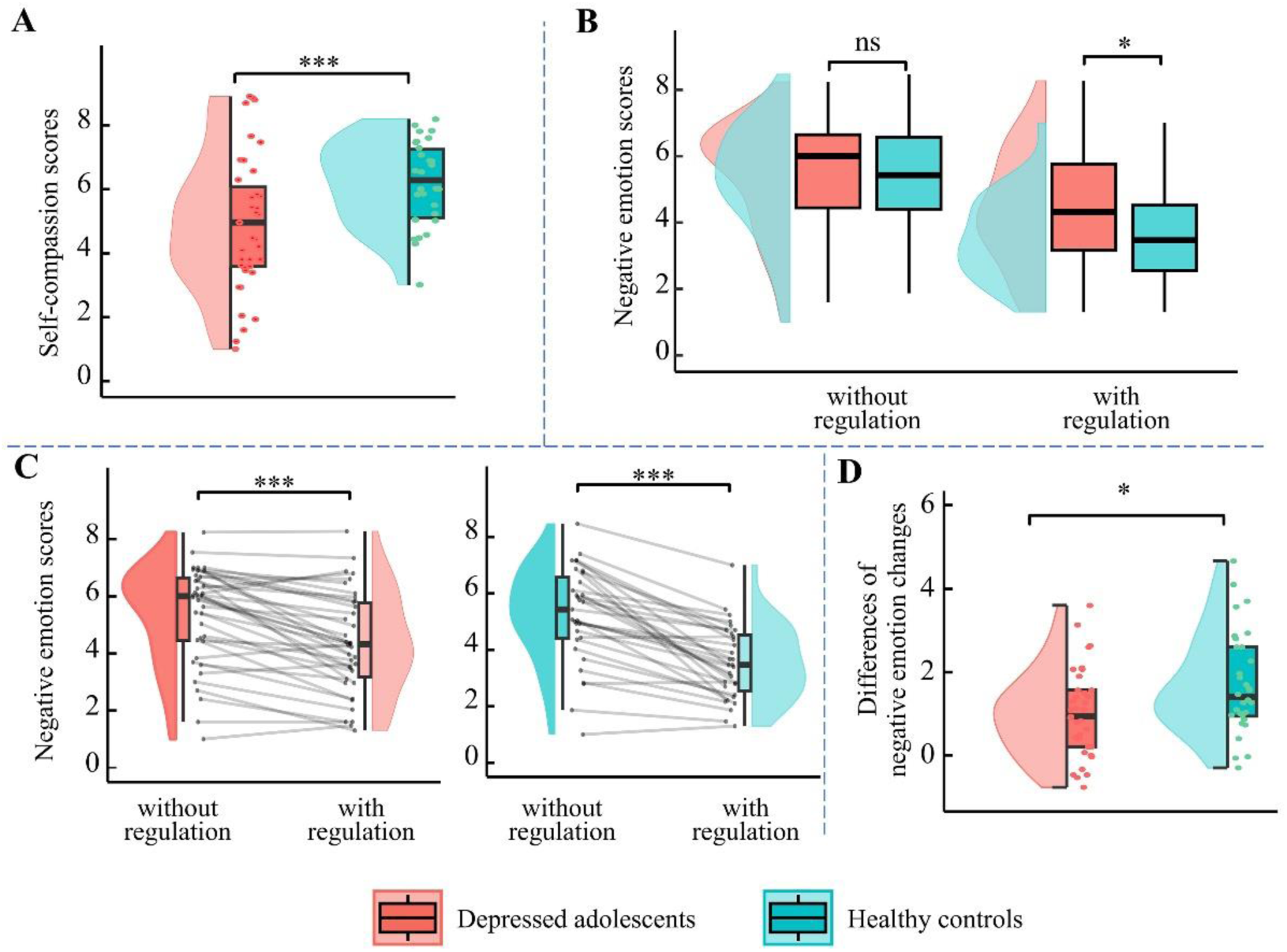
Behavioral results for self-compassion ratings and negative emotion across regulation conditions. (A) Self-compassion ratings during the regulation condition, showing group differences between healthy controls and adolescents with depression. (B) Negative emotion scores for both groups in the non-regulation and self-compassion regulation conditions. (C) Individual changes in negative emotion scores from the non-regulation to the regulation condition in adolescents with depression (left) and healthy controls (right). (D) Group differences in reductions of negative emotion (non-regulation minus regulation). Higher values indicate a greater decrease in negative emotion during self-compassion regulation.

**Figure 3.**
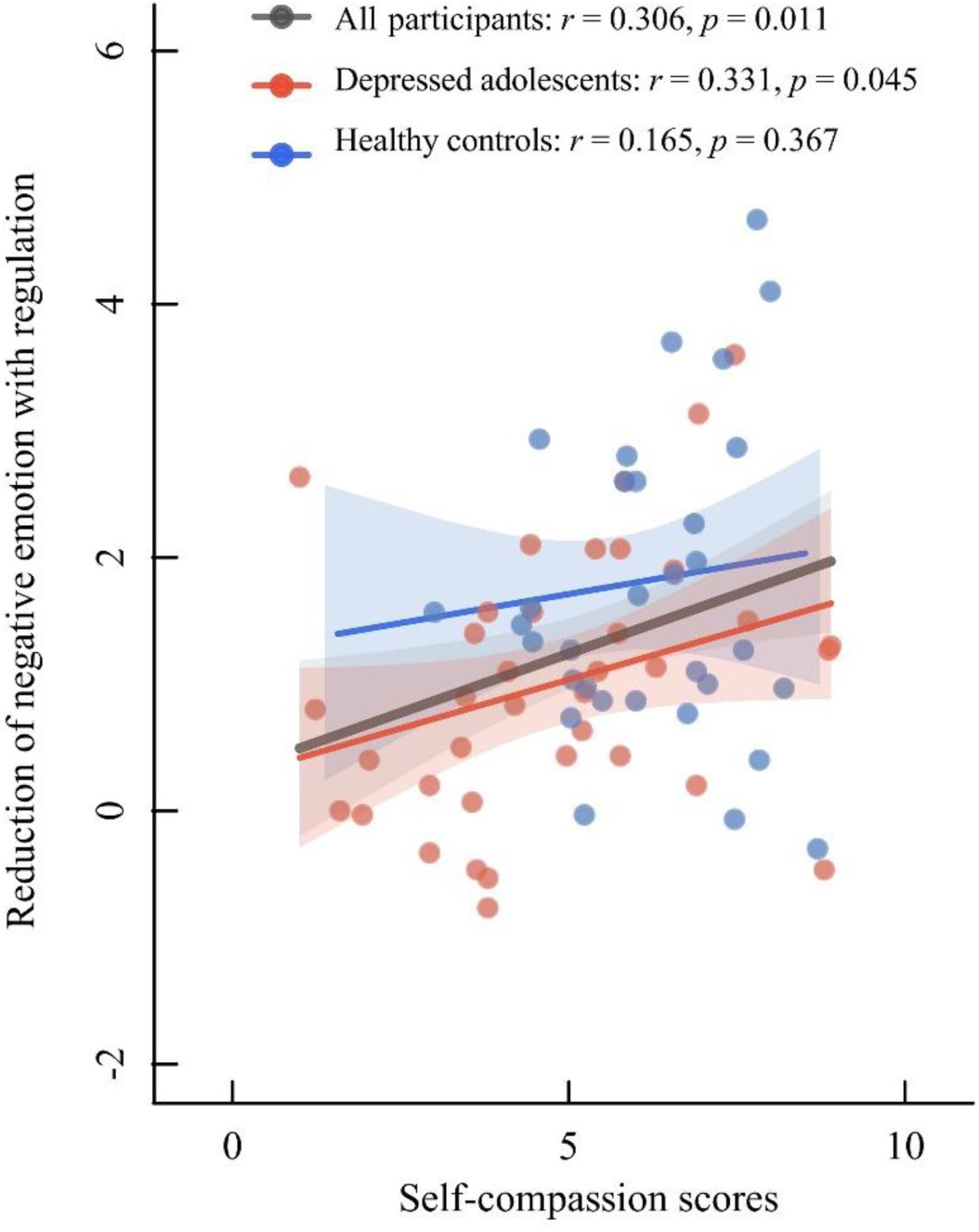
The partial correlations between self-compassion ratings and the reduction in negative emotion with regulation. These analyses were performed for all participants, the healthy controls, and adolescents with depression, controlling for age and sex.

In the stratified linear regression analysis, we selected the optimal model based on the model’s goodness-of-fit, shown as the differences of negative emotion scores = 0.35 + 0.14 × self-compassion ratings + 0.50 × group. In this model, both self-compassion ratings and group showed trends toward predicting the regulation effect (Table 2). Specifically, self-compassion ratings (*β* = 0.14, *t* = 1.81, *p* = 0.076) showed a marginally significant positive predictive effect on negative emotion scores, and group status (*β* = 0.50, *t* = 1.72, *p* = 0.090) also approached statistical significance, indicating that healthy controls exhibited greater improvement in negative emotion following regulation.

**Table 2.**
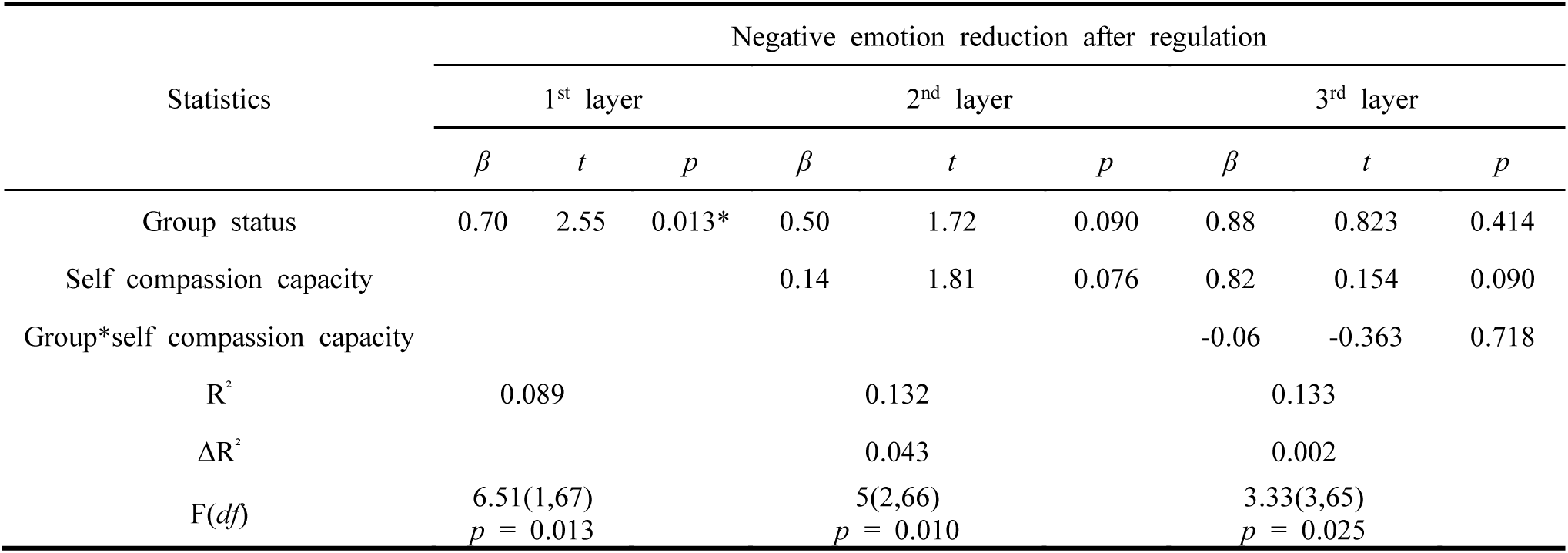
Stratified regression models predicting negative emotion reduction from self-compassion ratings.

### 3.2. Sustained high LPP response to social exclusion in adolescents with depression

ERP data revealed a consistent group difference in LPP during the task (Figure 4). The 2 × 2 repeated measures ANOVA observed no interaction effect between group and regulation conditions (*F*(1, 49) = 0.56, *p* = 0.459, *η*^2^ = 0.01), and no significant main effect of self-compassion regulation (*F*(1, 49) = 1.97, *p* = 0.167, *η*^2^ = 0.04). By contrast, there was significant main effect of group (*F*(1, 49) = 10.10, *p* = 0.003, *η*^2^ = 0.17), indicating sustained hyper-reactivity in depressed adolescents. Independent samples t-tests showed that the depressed group exhibited significantly larger LPP amplitudes compared to the healthy control group, both in the presence (*t* = 2.68, *p* = 0.011) and absence (*t* = 3.10, *p* = 0.003) of self-compassion regulation. Higher LPP amplitudes were associated with reduced changes in negative emotion ratings in healthy controls (*r* = -0.414, *p* = 0.044) but not adolescents with depression (*r* = 0.163, *p* = 0.457).

**Figure 4.**
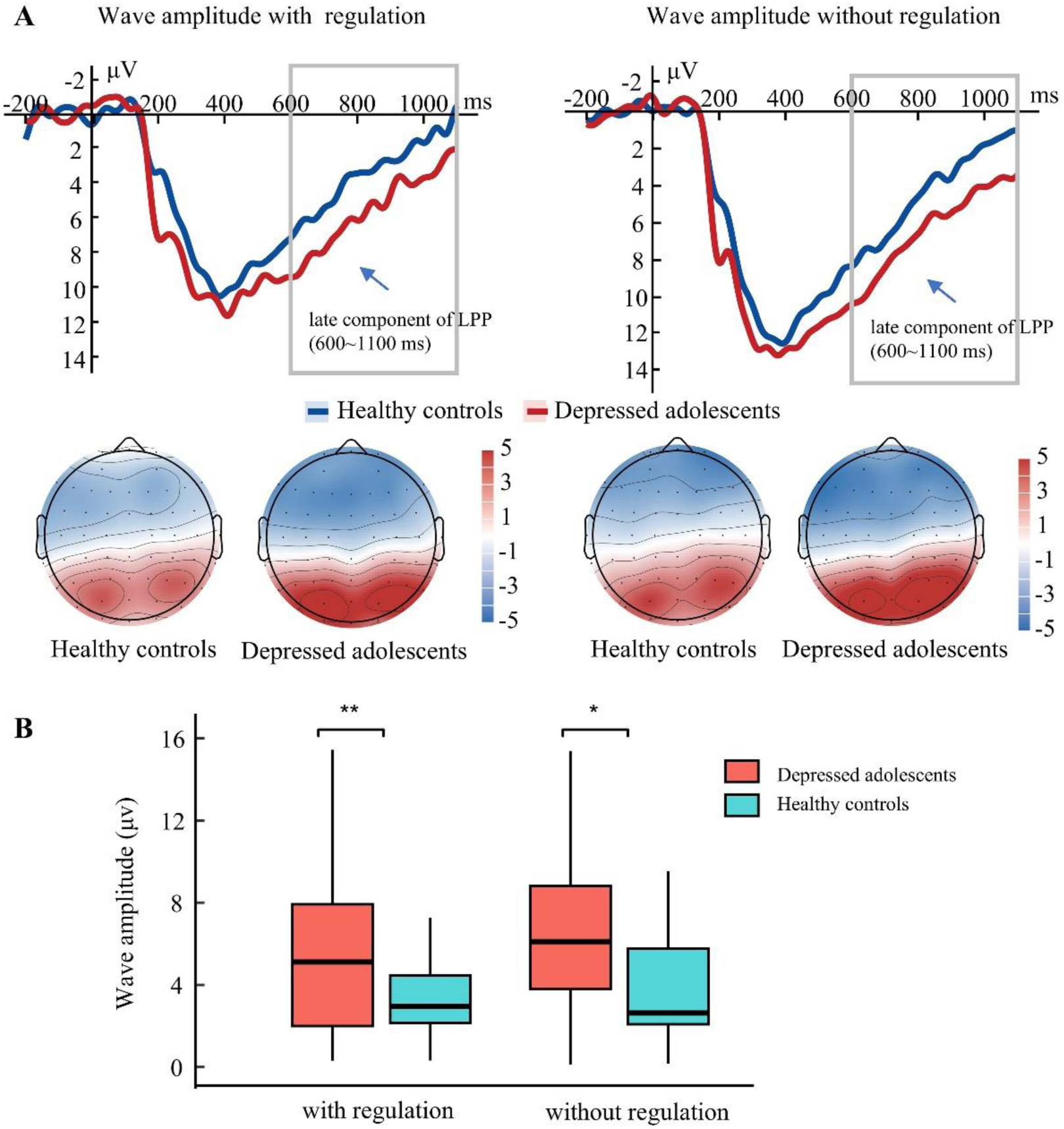
Late LPP component during self-compassion regulation in depressed and healthy control groups. (A) Grand-average ERP waveforms at selected electrodes PO3, PO4, and POz for each group, under the conditions with and without self-compassion regulation. The topographic maps displayed the scalp distribution of the late LPP component (600–1100 ms). The shaded gray regions denoted the time windows used for amplitude quantification. (B) Boxplots summarizing the repeated-measures ANOVA results for LPP amplitudes in each condition and group. The x-axis represented the regulation condition, the y-axis showed LPP amplitude, and group differences were indicated by separate lines.

## 4. Discussion

In this study, we examined the neural mechanisms underlying self-compassion regulation during social exclusion situations in adolescents with depression. Our findings demonstrated that adolescents with depression exhibited greater negative emotion, reduced engagement in self-compassion regulation, diminished emotion regulation effects, as well as a larger late LPP component. These findings suggest that negative emotional responses to social exclusion in adolescents with depression are characterized by heightened emotional reactivity alongside diminished effects of adaptive regulatory strategies such as self-compassion.

Our behavioral results showed that both healthy and depressed adolescents experienced negative emotions when confronted with social exclusion. Self-compassion reduced negative emotion ratings, which was consistent with previous findings [25, 26]. Compared with healthy controls, adolescents with depression demonstrated a significantly smaller reduction in negative emotion, which was related to their reduced engagement in self-compassion. Such functional deficits in negative emotion regulation in depressed individuals can be attributed to the excessive self-criticism and a lack of self-forgiveness and acceptance, which may exacerbate the negative emotional cycle [49]. Moreover, depressed individuals tended to engage in negative self-evaluation, and to hold beliefs such as being unworthy of love and being not good enough. These perceptions hindered effective emotion regulation and may make them more prone to experience stronger negative emotions when facing social exclusion [49, 50].

ERP results revealed that depressed adolescents showed higher LPP amplitudes during both experimental conditions compared with healthy controls. Taken together with the reduced emotion regulation effects, this may reflect potentially altered emotional processing during self-compassion regulation. The LPP is considered to reflect sustained attention and elaborative processing of emotionally salient stimuli [51, 52]. The LPP component is involved in emotional interpretive processing, and the magnitude of the wave typically indicates the depth of emotional stimulus processing and the degree of emotional regulation [53]. The enhanced LPP amplitude in adolescents with depression during social exclusion may reflect sustained, intensified processing of socially threatening affective information, suggesting difficulties in modulating responses to negative social experiences.

These findings may provide insights in interventions targeting emotion regulation difficulties in adolescents with depression. Previous studies suggested that self-compassion-based interventions may reduce depressive symptoms and alleviate emotional difficulties [54, 55]. Recent evidence has also demonstrated their potential efficacy in depressed adolescents from China in a 6-week self-compassion group psychotherapy programme [56]. Such interventions were generally low-cost, non-invasive, and can be delivered in clinical or school settings, making them potentially accessible and feasible for adolescents, thus representing a promising approach for adolescents with depression. However, further research is needed to determine the feasibility, accessibility, acceptability, adherence, and long-term efficacy with larger samples, as well as their underlying neurobiological mechanisms in adolescents with depression. In addition, our results suggested that brain stimulation interventions targeting the reduction of physiological responses may represent a complementary or potentially effective therapeutic approach alongside self-compassion-based interventions in adolescents with depression.

Our study had some limitations. First, our sample largely consisted of female adolescents. This pattern was consistent with the higher rates of depression among adolescent females than males [57], which has been attributed to multiple interacting factors such as pubertal development, biological vulnerability, and differential exposure to psychosocial stressors [58]. Although we included statistically sex-matched groups and controlled for sex as a covariate, the higher proportion of females and slightly unbalanced sex distribution may bias our findings. Given sex differences in emotion regulation processes and neural responses to emotional stimuli, including LPP amplitudes [59, 60], future studies with larger and more balanced samples are needed to improve generalizability.

Another limitation concerns the ecological validity of the social exclusion paradigm. Although the scenario imagination task allowed assessment of emotional responses to social exclusion while minimizing ethical concerns associated with experimentally inducing real interpersonal rejection, imagined scenarios may not fully capture the complexity and immediacy of real-life exclusion experiences. Previous studies have used more interactive social exclusion paradigms such as the Cyberball approach in children and adolescents, which may better approximate real-world ostracism-related distress [61]. Future studies could consider developmentally appropriate and ethically acceptable approaches, such as combining Cyberball paradigms and virtual reality, to further enhance ecological validity.

A further limitation concerns the temporal window of ERP analysis. In the present study, social exclusion-related images were presented for 8 seconds, whereas LPP analysis focused on the 600–1100 ms interval following stimulus onset. This approach was selected based on previous ERP studies examining sustained emotional processing during the late LPP time window [47]; however, this relatively early window may primarily reflect relatively early stages of emotional responses to social exclusion rather than the dynamic neural processes involved throughout the entire self-compassion regulation period. Future studies using time-resolved ERP analyses or other neuroimaging approaches could further characterize the temporal evolution of self-compassion regulation.

In addition, the cross-sectional design of our study limits causal inferences about the relationship between self-compassion regulation and depression. Longitudinal studies tracking the development of self-compassion and its relationship with depressive symptoms could provide valuable insights into whether impaired self-compassion is a risk factor, consequence, or maintainer of depression in adolescents.

Lastly, as the non-regulation condition did not involve an active comparison strategy, the observed reduction in negative emotion may partly reflect general cognitive engagement or attentional processes associated with applying self-compassion statements. Future studies using active control conditions are needed to further isolate the specific effects of self-compassion regulation.

## 5. Conclusion

Adolescents with depression exhibited deficits in using self-compassion to downregulate negative emotions, alongside sustained electrophysiological reactivity in response to socially threatening stimuli. These findings highlight an urgent need to develop interventions targeting brain activity that strengthen adolescents’ capacity to effectively deploy emotion regulation strategies such as self-compassion, thereby attenuating negative emotional responses in depression.

## Data availability

The data that support the findings of this study are available on request from the corresponding author, Kangcheng Wang.

## Author contributions

**Huijing Sun:** Conceptualization, study design, formal analysis and writing. **Wenjing Zou:** Conceptualization, study design, data collection, formal analysis and writing. **Duanwei Wang:** Conceptualization and data collection. **Yihao Zhang:** Data collection and study design. **Changlin Bai:** Data collection and study design. **Wenfu Li:** Data collection and study design. **Yanan Xiao:** writing. **Xiuqing Niu:** data collection. **Xuexiao Shao:** formal analysis. **Xiaoyue Wang:** writing. **Bernhard Hommel:** Conceptualization and study design. **Ying Yang:** Conceptualization, study design and data collection. **Kangcheng Wang:** Conceptualization, study design, data collection, formal analysis and writing.

## Declaration of generative AI and AI-assisted technologies

During the preparation of this work, we have used ChatGPT and DOUBAO to assist in drafting the scripts for data analysis. All AI-generated codes were carefully reviewed, verified, and edited by us. The authors declared that no AI tools were employed for writing the manuscript, interpretation of results or drawing the conclusions.

## Acknowledgements

None.

## Consent for publication

Not applicable.

